# Resurrecting ancestral genes in bacteria to interpret ancient biosignatures

**DOI:** 10.1101/164038

**Authors:** Betul Kacar, Lionel Guy, Eric Smith, John Baross

## Abstract

Two datasets, the geologic record and the genetic content of extant organisms, provide complementary insights into the history of how key molecular components have shaped or driven global environmental and macroevolutionary trends. Changes in global physicochemical modes over time are thought to be a consistent feature of this relationship between Earth and life, as life is thought to have been optimizing protein functions for the entirety of its ∼3.8 billion years of history on Earth. Organismal survival depends on how well critical genetic and metabolic components can adapt to their environments, reflecting an ability to optimize efficiently to changing conditions. The geologic record provides an array of biologically independent indicators of macroscale atmospheric and oceanic composition, but provides little in the way of the exact behavior of the molecular components that influenced the compositions of these reservoirs. By reconstructing sequences of proteins that might have been present in ancient organisms, we can identify a subset of possible sequences that may have been optimized to these ancient environmental conditions. How can extant life be used to reconstruct ancestral phenotypes? Configurations of ancient sequences can be inferred from the diversity of extant sequences, and then resurrected in the lab to ascertain their biochemical attributes. One way to augment sequence-based, single-gene methods to obtain a richer and more reliable picture of the deep past, is to resurrect inferred ancestral protein sequences in living organisms, where their phenotypes can be exposed in a complex molecular-systems context, and to then link consequences of those phenotypes to biosignatures that were preserved in the independent historical repository of the geological record. As a first-step beyond single molecule reconstruction to the study of functional molecular systems, we present here the ancestral sequence reconstruction of the beta-carbonic anhydrase protein. We assess how carbonic anhydrase proteins meet our selection criteria for reconstructing ancient biosignatures in the lab, which we term paleophenotype reconstruction.

## Main Text

The history of life on Earth has left two main repositories of evidence from which we can try to reconstruct it: the geological record, and the extant genetic diversity of organisms. Contemporary organisms, however, can be complex and cryptic vehicles for information about their history [1, 2]. Many genetic signatures in known life have been largely overwritten due to changing conditions of natural selection, evolutionary convergences, or simply genetic drift [3, 4]. However, functional links between the evolution of a protein (or a metabolic network) and biosignatures preserved in the geological record may provide clues to the timing and origin of major phylogenetic groups of organisms, including clades that went extinct and are no longer accessible for direct comparative study [5-9].

One way to connect the geological and genomic data sets is through ancestral sequence reconstruction [10-12]. One first infers ancestral sequences of biological molecules by phylogenetic reconstruction methods, and then uses these proposed sequences to synthesize models of paleoenzymes either computationally or experimentally [13-16]. In some cases the enzymes may be used to replace their modern counterparts in living organisms, being brought back to life as ‘revenant genes’, to obtain in situ expressions of “paleo-*phenotypes*” of the organisms that once harboured them [17, 18].

Efforts to reconstruct paleophenotypes require careful design, to avoid misinterpreting artefacts of reconstruction bias, or using host organisms that may be not faithfully reproduce ancestral phenotypes because too many of their systems have since adapted to other conditions. We present here a paleophenotype reconstruction approach that builds on prior efforts in paleoenzymology, extending the utilization of inferred ancestral gene / enzyme sequences engineered within modern organisms. Our functional framework builds on applying paleophenotypes to complex biology and on experimentally testing historical geobiological models and hypotheses. We begin by outlining the logical motivation for paleophenotype reconstruction and describe the criteria that should be addressed as a basis for selecting an enzymatic system for paleophenotype reconstruction at the systems level. We then use ancestral sequence reconstruction to determine the evolutionary history of a critical component of the photosynthetic CO_2_ fixation pathway – the beta-carbonic anhdyrase protein – and critically evaluate the selection criteria for candidate revenant genes suitable for paleopheotype reconstruction studies.

Our three linked goals are: 1) to learn (by solving concrete cases) when one must look beyond single-gene phylogeny to reconstruct entire *functioning molecular systems,* in order to correctly link enzyme properties to geological signatures; 2) by studying cases such as Calvin-cycle carbon fixation, where an isotope signature is inherently linked to a functional criterion such as molecular selectivity and an environmental property such as oxygen activity, to demonstrate a consistent multi-factor reconstruction of an organism’s phenotype in its environmental context; and 3) to search for features in which ancient proteins may have been truly more primitive than any proteins that have survived in extant organisms, to understand the evolutionary progression from the first (we may suppose fitful) invasions of new modes of cellular life, metabolism, or bioenergetics, and the refined forms of modern organisms that have made it difficult to infer the paths through which these emergences could have taken place.

### THE LOGICAL MOTIVATION FOR PALEOPHENOTYPE RECONSTRUCTION

To understand why – and for which systems – experimental paleophenotype reconstruction is likely to be an important scientific advance, it is helpful to reflect on the limited forms of information about evolutionary processes that are actually employed in conventional methods of sequence-based phylogenetic reconstruction. Efforts to map out more of the genotype-phenotype correspondence, whether through modelling or by *in vivo* expression, and to correlate these with independent evidence carried geologically, may be understood as a way to bring in other dimensions of information about evolution that can contribute to historical reconstruction.

#### Phenotype Information Can Augment Relatively Simple Sequence Substitution Models

The field of phylogenetic inference, after nearly a half century of dedicated work, has addressed most problems of consistent sampling and error estimation [19-27]. Yet the probability models that are the workhorses of most phylogenetic inference are disconnected from their context: they typically are site-local insertion, deletion, and substitution models with no semantics of the functioning objects produced.^1^ Often it is necessary to restrict probability models to evaluating substitution events independently at each site, to keep computations affordable especially for large data sets. However, such models are by construction incapable of reflecting interaction properties that can cause some joint variations to have very different likelihood to produce viable organisms than the marginal variations do independently.

Information about interaction effects can often be obtained from models of protein domain structure, folding, and function [30-37]. Learning how to use functional protein models to identify the most important non-local interactions and represent their effects on viability and fitness is one goal that can be pursued as more ancestral enzymes are reconstructed. Substitution probabilities must also be estimated jointly with alignments, and systematically biased alignment estimates can lead to mis-specified substitution models [38-41]. Information about folding and function can be particularly informative for ambiguous alignments, as substitutions or crossovers that preserve domain structures should yield viable organisms more often than those that would be incompatible with maintaining functional domains.

#### Synthetic-Biology Methods Offer Ways to Test the Internal Consistency of Reconstructions

Much current-generation phylogenetic inference, because of the big-data survey nature of its questions [42], yields independently-derived proposals for the presence or absence of genes in ancient genomes, along with putative sequences for ancient proteins [43-45]. However, the probability models generating these claims at present include no information (as part of the Monte Carlo generate-and-test cycle itself) about the consistency of the physiologies they predict. The use of synthetic biology methods to insert reconstructed genes into living organisms, or to test proposed molecular systems either *in vitro* or with modified genomes *in vivo* provides ways to test historical models at the system level. It can help bridge the gap between ancestral sequences inferred with algorithmically sophisticated but information-poor probability methods, and proposals for how they might have co-occurred in ancient cells.

### PROPOSED FRAMEWORK FOR REBUILDING PALEOPHENOTYPES IN THE LABORATORY

We outline here three criteria in choosing enzyme systems for which a paleophenotype reconstruction and systems-engineering approach may be feasible and may yield interesting insights beyond those delivered by simple sequence-phylogenetic methods alone. They are directed both at properties of the enzymes and properties of the clades and environments in which these occurred over time (see Figure 1).

**Figure 1.**
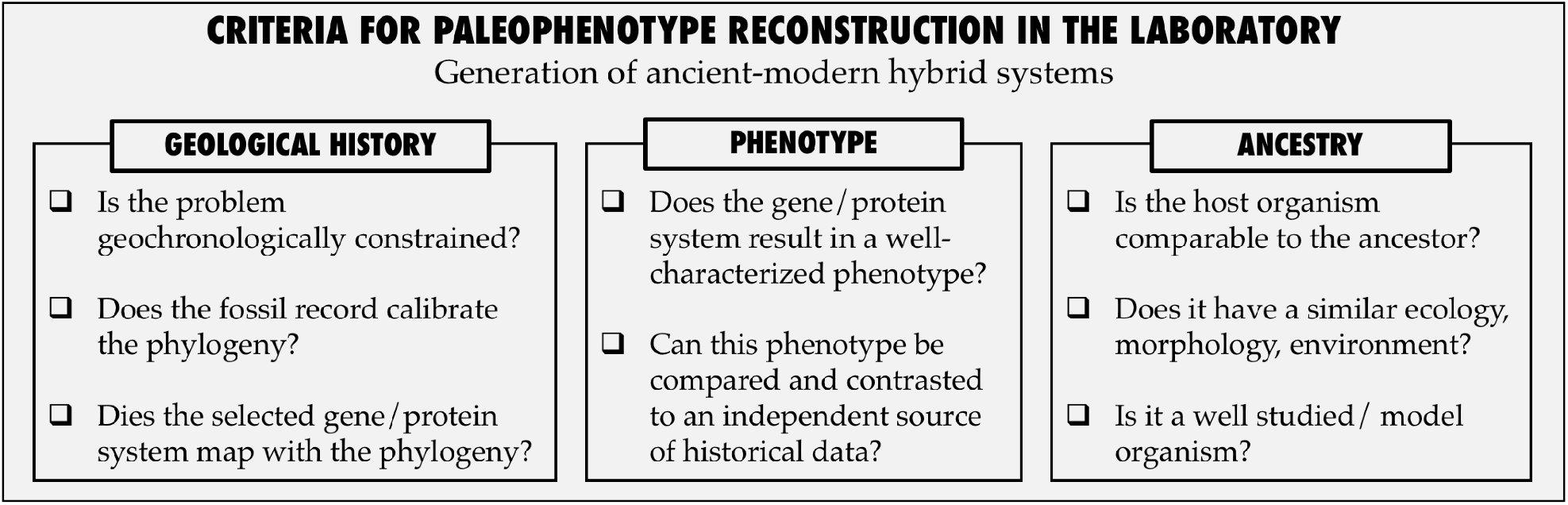
Criteria for paleophenotype reconstruction in the laboratory by generating hybrid ancient-modern bacterial systems

i. **Geology:** Is the problem geochronologically constrained? Does the protein system of interest mediate a biosignature that is recoverable from the rock record? Is there temporal structure in that biosignature that can be correlated with important evolutionary transitions in either enzyme context or function? Do major changes in enzyme function correspond to events of phylogenetic divergence, which can then be calibrated against geochronology?
ii. **Phenotype:** Can information be provided by *in vivo* resurrection of the protein that resolves important ambiguities in the usual methods of ancestral sequence inference, or that shows important errors in the assumptions usually made about sequence inference? This is information we think of as being reported by the phenotype of the protein, whether it is revealed by resurrection or by computational modelling.
iii. **Ancestry:** Do we have a current organism that is similar enough to the host organisms for the ancestral proteins that expressing them in our current organism will reveal the phenotypic characters that governed their function in the past? Is the proposed host a well-studied model organism? Are other essential components of metabolic pathways present in contemporary organisms also remnants of ancient life [46], and can their major evolutionary innovations also be inferred where these are significant to system functions?

Through reconstructing and examining the evolutionary history of contemporary components and then tying their phenotypes into biosignatures in mineral form, we can provide insight into innovations that are grounded in the rock record and thus in the geological and ecological context.

Enzymes involved in carbon-fixation such as Ribulose-1,5-bisphosphate carboxylase / oxygenase (Rubisco) proteins are thought to be one of the main causes of a distinct biosignature preserved in the rock record. This biosignature is revealed through comparison of 13C-isotope measurements between carbonate originally derived from atmospheric CO_2_ and organic carbon sequestered from biomass. 13C-isotope fractionation differences are the oldest record of living organisms, extending to at least ∼3.5 billion years in the past [47-50]. Rubisco is the distinctive catalyst and putative isotopic bottleneck for the Calvin-Benson-Bassham cycle [51, 52] – the predominant photosynthetic Carbon-fixation pathway by volume – and is therefore at the heart of many fundamental questions about the co-evolution of early life and the development of biogeochemical cycles of the planet [53]. Indeed, while we do not know whether ancient Rubisco proteins exhibited paleophenotypic properties that are comparable to those produced by contemporary Rubisco, or how efficiently the ancestral Rubisco proteins functioned under ancient environmental conditions, Rubisco plays a pivotal role in biogeochemical interpretation of the C-isotope fractionation patterns in deep time [47-50, 54-56]. Undoubtedly, characterization of ancient Rubisco as a means of elucidating steps of biochemical adaptation and resulting protein biochemical and organismal behaviour at the key nodes of phylogeny would be crucial and would be applicable for a paleophenotype reconstruction approach, suitable for subsequent isotope fractionation measurement and even phenotypic resurrection of isotope fractionation through engineering these ancient genes inside modern cyanobacteria [57-59].

Rubisco proteins don’t function in isolation in a cellular system. In bacteria, carbonic anhydrase proteins support Rubisco activity, by mediating efficient CO_2_ transport into and around the cell [60]. Carbonic anhydrase converts bicarbonate to carbon dioxide in the carboxysomes where Rubisco is localized – organelles thought to have evolved as a consequence of the increase in atmospheric oxygen concentration in the ancient Earth [61-64] – thereby alleviating the stringency required of Rubisco for carboxylase over oxygenase activity and reducing the energy and carbon loss that result from photorespiration. Additionally, carbonic anhydrases have essential roles in facilitating the transport of carbon dioxide and protons in the intracellular space, across biological membranes and in the layers of the extracellular space [65].

To understand the later-diverging innovations of photosynthetic systems associated with the rise of oxygen, the drawdown of atmospheric and oceanic CO_2_, and the colonization of land, it may even become essential to jointly reconstruct innovations in Rubisco and carbonic anhydrase with other metabolic and compartmental systems that serve as Carbon-concentrating mechanisms [66, 67]. The joint evolution of pathways associated with photorespiration may also provide evidence about O_2_/CO_2_ discriminatory capabilities of ancestral enzymes as well as the interpretation of the ancient isotope signals.

As the first step beyond single-molecule reconstruction to the study of functional molecular systems, we present here the phylogenetic history of carbonic anhydrase enzymes and the ancestral sequence for the beta-carbonic anhydrase protein. We assess whether/how carbonic anhydrase proteins meet our selection criteria for paleophenotype reconstruction, and demonstrate that events of horizontal gene transfer in an evolutionary tree for a given gene need to be recognized prior to a laboratory paleophenotype reconstruction.

### A UNIVERSAL CARBON SHUTTLE IN PHOTOSYNTHESIS AND BEYOND: CASE STUDY OF THE RECONSTRUCTION OF ANCIENT CARBONIC ANHYDRASE PROTEINS

Carbonic anhydrase is found in metabolically diverse species representing all three domains of life, [68-70]. The three main classes of carbonic anhydrase (alpha, beta and gamma) are not homologous and are thought to be a result of convergent evolution [71, 72]. Although molecular dates based only on sequence comparison should be regarded with caution, it has been suggested that both the gamma and the beta classes are ancient enzymes, which existed before the split between archaeal and bacterial domains [73, 74].

In a coarse assessment, carbonic anhydrase meets our paleophenotype selection criteria. It carries out an essential and ancient function in the carbon concentration machinery. While no particular study (to our knowledge) has attributed a specific biosignature to the activity of bacterial carbonic anhydrases, this enzyme mediates CO_2_ efflux in the carboxysome, potentially impacting the interpretation of Rubisco kinetic isotope selectivity, which is correlated with molecular CO_2_/O_2_ discrimination and turnover rate, in terms of the ambient CO_2_ and O_2_ activities in the cellular environment. The root of the gamma-class is inferred to have extended to approximately 4.2 billion years ago [73]. Moreover, the presence of carbonic anhydrase in thermophilic chemolithoautotrophs suggests that other ancient CO_2_-fixation pathways besides the Calvin cycle also depended on carbonic anhydrase function for efficient C-fixation [75, 76].

In this study, we focus on the beta-carbonic anhydrase, also called the prokaryotic carbonic anhydrase (although it has been found in eukaryotes as well). Beta-carbonic anhydrase is an ancient enzyme, it is widely represented in prokaryotes and despite its critical role for Earth’s biosphere, to date, not many studies focused on the molecular evolution of beta-carbonic anhydrases [77, 78]. Beta-carbonic anhydrases have been subdivided into four main clades, A to D. One group of enzymes belonging to the B clade of the beta-carbonic anhydrase is a probable example of neofunctionalization: these enzymes take CS2 (and not CO_2_) as a substrate [79].

The alignment of the 457 beta-carbonic anhydrase proteins revealed a strong conservation of about 200 amino-acids, of which a handful are identical in all or almost all sequences. The phylogenetic reconstruction of the carbonic anhydrase from 388 genomes confirms the existence of four, relatively well-supported clades (Figure 2 and Supp. Figure 1). Clade D seems to be the most distant one, and based on this we have chosen to root the tree from this group. The presence of archaeal homologs close to the root of three out of the four clades gives some credence to the hypothesis that duplication of the beta-carbonic anhydrase is very ancient, perhaps occurring before the archaea/bacteria separation. Except for the main clades and a few other shallower groups, the bootstrap values are altogether low, an ambiguity often observed when reconstructing deep phylogenies based on short protein alignments, due to small amount of genetic information available [80]. This is further emphasized by the fact that the main bacterial phyla – with the notable exception of cyanobacteria – were rarely reconstructed as monophyletic, even inside each clade. The incongruence between the carbonic anhydrase tree and the generally accepted tree of life can be explained by either one, or a combination of two, main factors: (a) the amount of phylogenetic information contained in this 200-site alignment might be too low to reliably approximate the true tree, or (b) the carbonic anhydrase has been horizontally transferred several times. Amongst the few clades, the two that are almost solely comprised of sequences from the same phylum are two groups of cyanobacteria, one belonging to clade B (bootstrap support: 98) and one to clade C (low bootstrap support) (Figure 2). In addition, sequences from cyanobacteria are found almost exclusively in these two clades. Despite the poor support values of the tree, even inside the group encompassing the cyanobacterial sequences of clade B, a likely scenario for the evolution of the carbonic anhydrase in cyanobacteria can be drafted. We hypothesize that the last common cyanobacterial ancestor had at least two copies of the gene, one clade B-like and the other clade C-like, and that one or the other copy was lost (and perhaps further copies gained) following different patterns in different cyanobacterial descendants. In consequence, we performed an ancestral reconstruction only for the well supported, monophyletic cyanobacterial group in clade B (see Supp. Figure 2).

**Figure 2.**
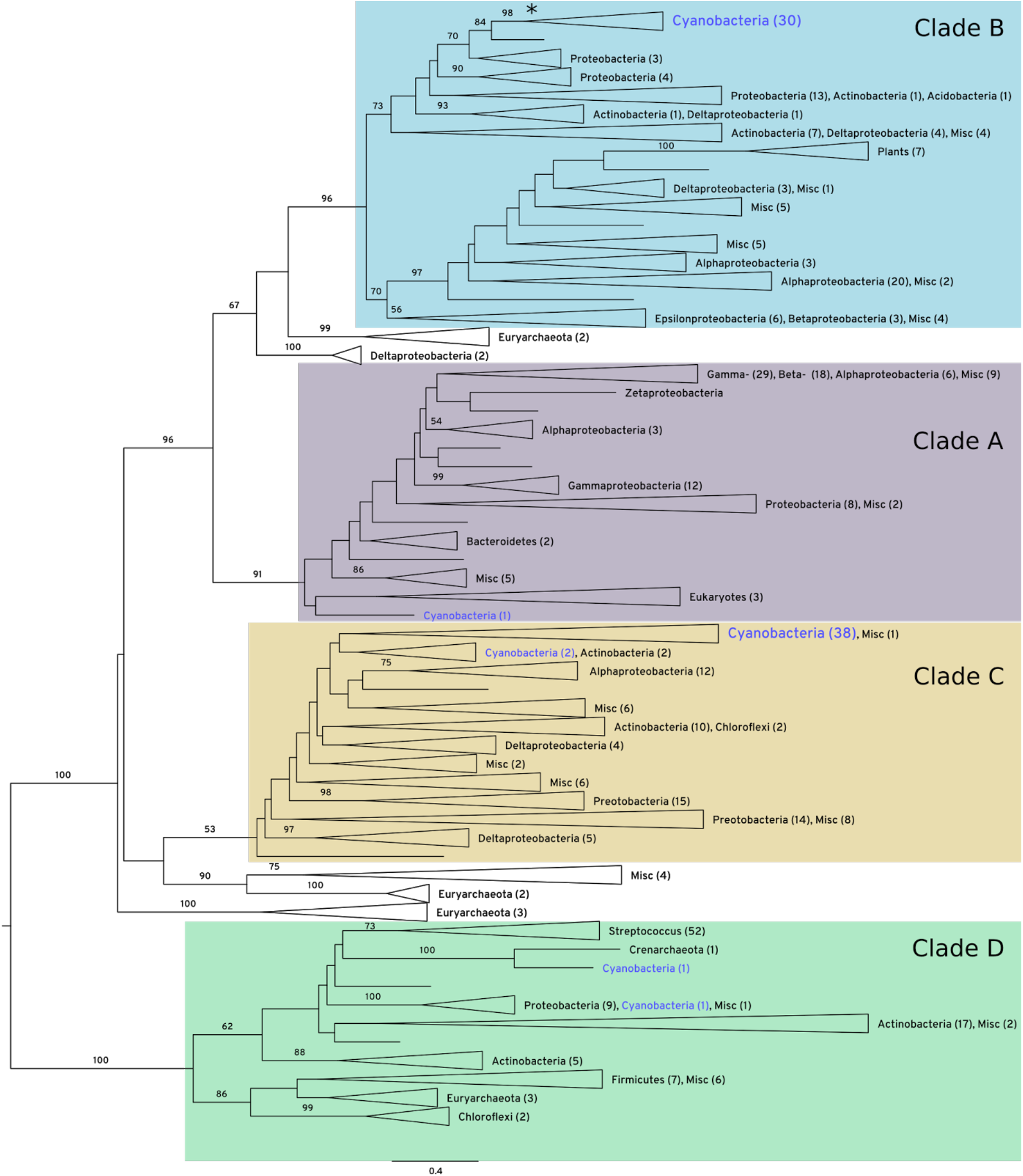
Maximum-likelihood phylogenetic reconstruction of the carbonic anhydrase (CA). The 492 CA homologs retrieved from 388 representative genomes were aligned with mafft-linsi and a maximum-likelihood phylogeny was inferred from the alignment with RAxML, using the PROTCATLG model. Bootstrap support is displayed for branches supported by > 50 bootstrap trees. Clades were collapsed to provide a more readable tree. The number of members of major taxonomic groups is presented in parentheses next to each collapsed clade. Cyanobacteria are depicted in blue. The four major recognized clades of CA (A-D) are highlighted in different colors. An asterisk marks the localization of the ancestor of the cyanobacterial clade B carbonic anhydrase ancestor. The scale at the bottom of the tree represents the number of substitution per site. The complete tree is depicted in Supp. Figure 1.

Previously reconstructed phylogenetic history of the Rubisco proteins, an essential partner of carbonic anhydrase in the carboxysome, displays a highly supported phylogenetic tree, which recapitulates the organismal phylogeny [55, 81]. In contrast, here we show that the evolutionary history of the carbonic anhydrase is more complex: several enzymes with no common ancestor (the different classes of carbonic anhydrases) catalyse the same reaction. Even within classes of carbonic anhydrases, duplications and horizontal gene transfers seem frequent. This is an expected outcome of the greater generality and modular function of carbonic anhydrase compared to the specialized role of Rubisco: modular components of a metabolic system are much more readily transferred^2^ or re-evolved through convergent evolution. The interpretation that carbonic anhydrase function is general and modular is further suggested by its redundant presence in many organisms: the number of homologs of the beta-carbonic anhydrase ranged from none to as many as six carbonic anhydrase genes in cyanobacteria, with most genomes having more than one gene. This suggests that the cost of exchanging (by horizontal gene transfer) or losing one copy of carbonic anhydrase (either completely or by neofunctionalization, as in the case of the CS_2_ hydrolase [79] is small. Nevertheless, ancestral reconstruction, in parts of the tree where a phylogenetic signal can be established with significant confidence, as in the cyanobacterial group in the B clade of the beta-carbonic anhydrase, shows that the conserved parts of the alignment are even more conserved in the ancestors of the clade than in the extant species (Figure 3). None of the highly conserved residues differs in the ancestors of the group (Node 67, depicted with an asterisk on Figure 2, and its descendants, but not the extant species), suggesting that the function of the enzyme hasn’t significantly changed inside the group.

**Figure 3.**
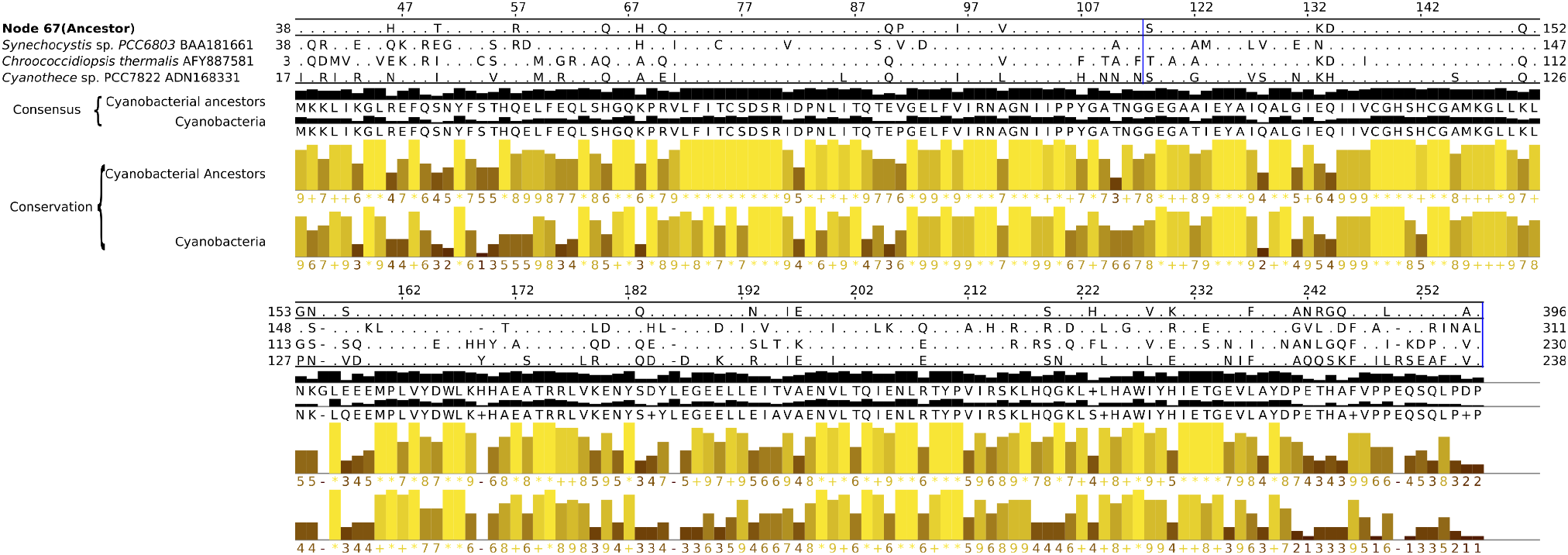
Alignment and analysis of ancestors and extant sequences of the cyanobacterial group in clade B beta-carbonic anhydrase. The alignment is based on all 55 ancestors and 57 extant sequences. From top to bottom: the aligned sequences for the last common ancestor of the cyanobacterial sequences (Node 67, shown with an asterisk in Figure 2), and three extant sequences. Only the amino acids different from their respective consensus sequences below (black bar graph) are shown. The yellow bar graphs represent the conservation in the two groups. Columns of the alignment that had more gaps than sequences are not represented. The raw alignment is available at http://phylobot.com/38899544/.

We further analysed the reliability of the ancestral carbonic anhydrase (Node 67) in the last cyanobacterial common ancestor by examining the posterior probabilities for each reconstructed residue (Figure 4). Multiple alignment of the reconstructed ancestor sequence with the sequences from all of the known extant species shows that the sequence of the large domain (located on the N-terminus side) can be confidently established (Figure 4). This fragment ranges from residues 40 to 240, which covers the functionally important zinc-binding core [83, 84]. On the other hand, the other parts of the reconstructed protein sequence are less reliable, as evidenced by the fact that the second- and third most probable amino-acid are closer to the most probable one (Figure 4).

**Figure 4.**
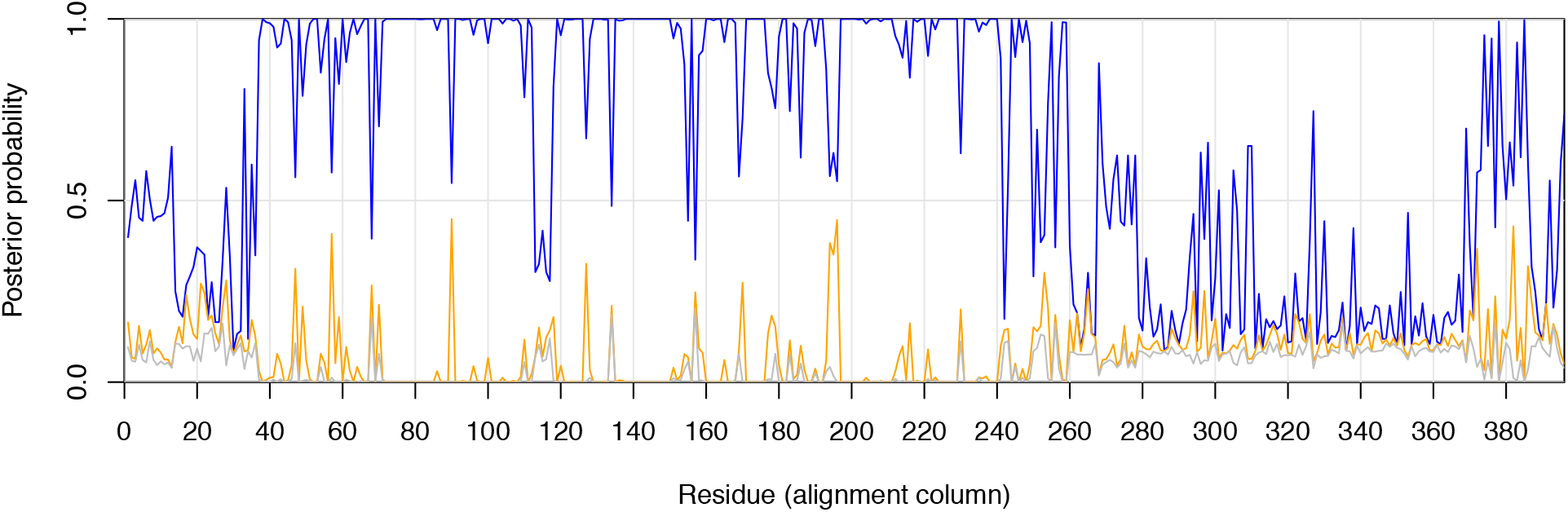
Posterior probabilities of maximum likelihood sequence residues, per position for the best (blue), second (orange) and third (grey) best residue, as reconstructed by phylobot.

### SIGNATURES OF PRIMITIVENESS: EXTENDING LESSONS LEARNED FROM PHYLOGENETIC RECONSTRUCTION TO MORE GENERAL PRINCIPLES OF MOLECULAR EVOLUTION

From joint phylogenetic/geochronological reconstructions such as our reconstruction of Rubisco and carbonic anhydrase, we may attempt to identify more general principles of molecular evolution, either by finding *signatures of primitiveness* – characters in which the first invaders of a new niche or new functionality still reflect what they were before, and are not yet well-adapted to their new mode of life – or by studying the dynamic by which forces of selection become *displaced from one. molecular system to another,* when some primitive, inherited molecular function is incapable of responding to selective criteria from the new environment that have become irresistible.

Relevant signatures of primitiveness can vary across protein families and ancestral functions. In some cases, it may be expected that proteins which are now sub-functionalized to specific substrates were once multifunctional, a change that we would expect to see [85-87] in a deep past when error rates in genome replication and also in translation should have been higher [88], favouring fewer and shorter genes, at the cost that each gene may have been required to catalyse multiple reactions in order for pathway formation to be possible.^3^

The very existence of carboxylase / oxygenase discrimination in Rubisco has the character of an emerging but stalled sub-functionalization. Molecular oxygen was by all evidence a minor component of the environments in which Rubisco emerged, and the enzyme mechanism was selected on the only relevant substrate: CO_2_. The ability of cyanobacteria to perform oxygenic photosynthesis is thought to have converted the early reducing atmosphere into an oxidizing one, which dramatically changed the composition of life forms on Earth by simultaneously enabling new life forms tolerant to oxygen and leading to the near-extinction of the existing ones [91, 92]. By enabling the massive proliferation of oxygenic photosynthesizers, Rubisco introduced the need for a substrate discrimination that had not existed when it arose, potentially creating conditions for its own failure. The reaction mechanism to which the whole enzyme structure is committed is one for which discrimination is costly and only partially successful, even under intense selection pressure. The result was *displacement* of selective pressure from Rubisco onto other enzymes such as carbonic anhydrase, and onto cellular ultrastructure in forming the carboxysome.

Due to the correlation between isotope selectivity and substrate discrimination in Rubisco, a further signature of primitiveness is suggested, which can readily be empirically tested. All modern Rubiscos fall along a rather tight linear regression between turnover rate and CO_2_/O_2_ discrimination, with a less-tightly correlated isotope shift [51, 52]. The apparently bright horizon, beyond which no Ribiscos are found, is the reason for the interpretation of an inherent trade-off in the mechanism that fixes CO_2_ using only the free energy of hydrolysis, which forces turnover to be sacrificed as the price of discrimination. The absence of dispersion on the low-performance side of the regression has been interpreted as evidence of evolutionary optimization: that turnover is always maximized against the futile cycle of photorespiration in the CO_2_/O_2_ environment of the enzyme. By studying turnover versus discrimination in ancestral Rubiscos, we can test whether they seem to reflect the same optimization horizon as modern Rubiscos. If not, one possibility is that the enzymes were more primitive; another is that naive sequence reconstruction methods miss essential information needed to identify the true ancestral form for this protein. Coupling an optimality analysis with functional measures of carbonic anhydrase will then allow us to compare the implied CO_2_/O_2_ environment to ambient conditions in eras suggested by the phylogenies of Rubisco and other molecular clocks or ancient biosignatures.

### EXTRAPOLATING BIOLOGY BACK TO EARLIEST- OR PRE-BIOLOGICAL CONDITIONS

Comparative sequence reconstruction, even augmented by geochronology, can only reveal directly historical evidence within the era from which sequence divergences have been preserved to the present. The geochronological evidence may extend to earlier times – as is potentially the case for organic carbon signatures, although these become sparse near the time when life emerged on Earth about 4 billion years ago [93-95] – and it is plausible that complex cellular life also existed and evolved within these eras; but to understand them we will need interpretive methods beyond simple sequence comparison.

Phylogenetic reconstruction should move beyond sequence reconstruction and the functional deductions based on phenotypes that are heavily impacted by the sequence reconstruction quality. Indeed, early examples in the young field of paleoenzymology attempted to make inferences about the temperature of ancient environments that no longer exist, by interpreting ancestral protein sequences from organisms long-since extinct [96-99]. Its predictions should come to integrate enzymatic structure and folding information as well as systems-level effects on protein network interactions and physiology – hence the emphasis on “paleophenotype” rather than “paleosequence” or “paleostructure”.

Engineering ancient pathways whose behaviour could recapitulate certain past phenotypes and innovations has the potential to reconcile ambiguities in phylogenetic reconstruction. This may be realized by focusing on assessing those particular phenotypes that facilitate effective comparison to an independent historical record of component or organismal phenotype contained in the rock record. The same dynamical effects – error-prone replication, horizontal gene transfer [88, 100-103]– that tend to erase memory about genes and genomes in the deepest eras of life, also remove one of the aspects of biology that interferes most with modelling from first principles: the capacity for historically-contingent features to contribute essential context for function. If we can use a combination of deep sequence reconstruction, functional modelling, and geological constraints on phenotype, we may be able to identify the “rules of assembly” for very early living systems. These are the rules that, as we extend to epochs in which memory was less robust, should have governed strong evolutionary convergences, and for the earliest molecular systems that were bound in the most detail to their geological environments, they may permit a limited amount of prediction from first principles about what those systems could have been. The reconstruction of carbonic anhydrase demonstrates the potential for integrative investigation. Though the carbonic anhydrase tree shows that horizontal gene transfer is a barrier for deep time reconstructions of some enzymes using single gene trees, carbonic anhydrase’s intimate interactive relationship with other enzymes of the carbon uptake and carbon-concentrating mechanism apparatus leaves open the possibility of isolating its functional behaviour in conjunction with enzymes with more clearly resolved genetic histories such as Rubisco.

### CONCLUSION

The ability of cyanobacteria to perform oxygenic photosynthesis is thought to have converted the early reducing atmosphere into an oxidizing one, which dramatically changed the composition of life forms on Earth by simultaneously enabling new life forms tolerant to oxygen and leading to the near-extinction of species acclimated to anoxic conditions. Much of this information is derived from the geologic record- evidence of carbon cycling (and biological activity) can be inferred from carbon isotopes which lies at the interfaces between enzymatic activity, organismal phenotype and the formation of sedimentary rocks [104]. As a corollary to these interpretive schemes, it is assumed that the controlled fixation of inorganic carbon to organic carbon is a precondition for the emergence of living systems. While selectivity in the carbon isotope composition of biologically produced organic matter is evidence that is preserved in the rock record that reflects the metabolic activity of ancient organisms, this record is increasingly problematic near the time when life emerged on Earth about 4 billion years ago. Means of investigating life’s early evolution and origins that complements the loss of a robust geologic record may prove insightful.

Characterizing paleophenotypes in the laboratory is now feasible with evolutionary techniques already available and new techniques currently under development. However, the best chance of succeeding is in the integration of ancestral sequence reconstruction, genetic engineering and laboratory evolution. Current efforts attempt to establish the extent to which biochemical properties of paleoenzymes may be correlated with biosignatures retrievable from the rock record, such as stable isotope ratios [57]. This behavioural information is critical to understanding ancient biological innovations and their effect upon (and shaping by) the Earth’s environment. It is likely that the specific genotype / phenotype relationship in present-day organisms has evolved from organisms with very different metabolisms. If so, when and where did specific extant phenotypes evolve? How conserved are the genes of interest through time? How frequently were these genes transferred to other organisms? What can paleophenotypes tell us about the origins of critical metabolic pathways? Reprogramming contemporary organisms by engineering their genomes with ancestral DNA is a fundamentally new methodology with which to extrapolate back to the origins of life. The technique is part of our broader approach that seeks to reconstruct macroevolutionary phenotypic trends across geologic time to ultimately infer conditions that approach the Last Universal Common Ancestor and possibly life’s origins.

## Methods

### Genome selection

Representative genomes for different taxonomic groups were selected using the software phyloSkeleton (manuscript in revision; available at https://bitbucket.org/lionelguy/phyloskeleton) as follows: one for each species of *Streptococcus,* one for each family in Proteobacteria and Cyanobacteria, and one for each order otherwise. After manual curation of poorly classified genomes, 388 genomes were retained (Supp. Table 1).

### Carbonic anhydrase homologs identification

Genomes were search with HMMer v3.1b2 [105], using the PFAM profile for the so-called prokaryotic carbonic anhydrase (http://pfam.xfam.org/family/PF00484), which contains representatives of the beta-carbonic anhydrase clades A, B and C (but not D), as defined in [79]. All homologs with an E-value < 1e-10 were retained. In the 388 selected genomes, between 0 and 6 homologs of CA per genome were found, with most genomes having 0 (102 genomes), 1 (176 genomes) or 2 (72 genomes) copies. In total, 457 sequences were retrieved. The sequence length varied from 118 to 867 amino acids, with eukaryotic homologs being the longest. Most sequences were between 200 and 250 amino acids.

### Sequence alignment

The 457 CA homologs found by phyloSkeleton and the 35 sequences that could be retrieved from Smeulders et al. (2011) were aligned with mafft-linsi v7.215 [106] and the resulting alignment were filtered to remove positions that had >50% gaps, using trimal [107]. The filtered alignment counted 200 positions and was visually inspected for any obvious misaligned regions.

### Phylogenetics

RAxML 8.2.8 [108] with the PROTCATLG model was used to infer the phylogenetic tree depicted in Figure 2. A hundred parametric bootstraps were drawn to estimate branch reliability. Trees were visualized and edited with FigTree 1.4.3 (Andrew Rambaut, available from http://tree.bio.ed.ac.uk/software/figtree/).

### Ancestral reconstruction

Ancestral reconstruction of the clade B carbonic anhydrase in cyanobacteria was done by first collecting all homologs of the carbonic anhydrase in cyanobacteria, choosing one representative genome per genus in cyanobacteria with phyloSkeleton, and adding genomes branching close to cyanobacteria, both clade A and clade B *(Sorangium cellulosum, Sandaracinus amyloliticus,* and *Beijerinckia indica)* in Figure 2, resulting in 83 genomes (Supp. Table 2). Sequences belonging to clade B were then extracted and uploaded to Phylobot [109]. Sequences were analyzed with muscle and msaprobs, and trees were drawn under the PROTCATLG and PROTGAMMALG models. The complete results are available at Phylobot: http://phylobot.com/38899544/. The result of the muscle alignment and the tree drawn under PROTCATLG were further analyzed and visualized in Jalview 2.10.1 [110].

## Authors’ Contributions

All authors analyzed the data, contributed to writing the final manuscript and gave final approval for publication.

## Supplementary Figures

Supplementary Figure 1: Maximum-likelihood phylogenetic reconstruction of the carbonic anhydrase. Same analysis as for Figure 2, but no clade has been collapsed, and all bootstrap values are shown. Leaves are coloured according to their taxonomic classification: red, Proteobacteria; blue, Cyanobacteria; green, Archaea; orange, Actinobacteria; purple, Firmicutes. When available, the organism name is prefixed with the phylum and class the organism belongs to.

Supplementary Figure 2: Maximum-likelihood phylogeny of carbonic anhydrase homologs present in representative of each genus of the cyanobacteria and a few outgroup species. The tree was inferred with RAxML. Bootstrap support is shown for each branch. Shades of blue represent different families of Cyanobacteria. The scale represents the number of substitutions per site.

## Supplementary Tables

Supplementary Table 1: List of 388 genomes initially searched for the presence of carbonic anhydrase. The file is a partial output of phyloSkeleton.

Supplementary Table 2: List of 79 cyanobacterial genomes and 4 outgroups. The file is a partial output of phyloSkeleton.

## Footnotes

1 Here we make a distinction that is standard in the information theory of symbol sequences, between *syntactic* information, derived from probability models for uninterpreted sequence strings like the stringsubstitution models used in molecular phylogenies, and *semantic* information, which informally we think of as deriving from functional interpretations of those strings, and which can sometimes be formalized as mutual information between the symbol sequences and their environments [28] Adami, C. 2002 Sequence complexity in Darwinian evolution. *Complexity* **8**, 59-56, [29] Adami, C. 2004 Information theory in molecular biology. *Physics of Life Reviews* **1**, 3-22.

2 Perhaps the most striking contrast between modular and non-modular molecular components in their degree of horizontal gene transfer is recognized between proteins and RNA associated with the ribosome, and the aminoacyl-tRNA synthases. The former are proxies for a reference Tree of Life, while the latter have been ubiquitously subject to waves of replacement [82] Woese, C. R., Olsen, G. J., Ibba, M. & Soll, D. 2000 Aminoacyl-tRNA synthetases, the genetic code, and the evolutionary process. *Microbiol Mol Biol Rev* **64**, 202-236.

3 Such an interpretation has been advanced for homologies in proteins of the rTCA cycle in Aquificaceae [89] Aoshima, M. & Igarashi, Y. 2008 Nondecarboxylating and decarboxylating isocitrate dehydrogenases: oxalosuccinate reductase as an ancestral form of isocitrate dehydrogenase. *J Bacteriol* **190**, 2050-2055. (DOI:10.1128/JB.01799-07), [90] Aoshima, M., Ishii, M. & Igarashi, Y. 2004 A novel enzyme, citryl-CoA synthetase, catalysing the first step of the citrate cleavage reaction in Hydrogenobacter thermophilus TK-6. *Mol Microbiol* **52**, 751-761. (DOI:10.1111/j.1365-2958.2004.04009.x)., an alternative carbon-fixation cycle likely predating the Calvin cycle.

## References

[1] Gould, S. J. 1989 Wonderful life. New York, NY, Ww Norton.

[2] Koonin, E. V. The logic of chance: the nature and origin of biological evolution. Upper Saddle River, NJ, FT Press.

[3] Dobzhansky, T. 1937 Genetics and the origin of species. New York, NY, Columbia University Press.

[4] Huxley, J. S. 1942 Evolution: the modern synthesis. London, UK., Allen and Unwin.

[5] Gogarten, J. P., Fournier, G. P. & Zhaxybayeva, O. 2008 Gene Transfer and the Reconstruction of Life’s Early History from Genomic Data. In Strategies of Life Detection (pp. 115–131.

[6] Goldford, J. E., Hartman, H., Smith, T. F. & Segre, D. 2017 Remnants of an Ancient Metabolism without Phosphate. Cell 168, 1126–1134 e1129. (DOI:10.1016/j.cell.2017.02.001).

[7] Martin, W. F. & Thauer, R. K. 2017 Energy in Ancient Metabolism. Cell 168, 953–955. (DOI:10.1016/j.cell.2017.02.032).

[8] Cornejo-Castillo, F. M., Cabello, A. M., Salazar, G., Sanchez-Baracaldo, P., Lima-Mendez, G., Hingamp, P., Alberti, A., Sunagawa, S., Bork, P., de Vargas, C., et al. 2016 Cyanobacterial symbionts diverged in the late Cretaceous towards lineage-specific nitrogen fixation factories in single-celled phytoplankton. Nat Commun, 7 11071. (DOI:10.1038/ncomms11071).

[9] Fischer, W. W. 2008 Biogeochemistry: Life before the rise of oxygen. Nature 455, 1051–1052. (DOI:10.1038/4551051a).

[10] Le, S. Q. & Gascuel, O. 2008 An improved general amino acid replacement matrix. Mol Biol Evol, 25 1307–1320. (DOI:10.1093/molbev/msn067).

[11] Perez-Jimenez, R., Ingles-Prieto, A., Zhao, Z. M., Sanchez-Romero, I., Alegre-Cebollada, J., Kosuri, P., Garcia-Manyes, S., Kappock, T. J., Tanokura, M., Holmgren, A., et al. 2011 Single-molecule paleoenzymology probes the chemistry of resurrected enzymes. Nat Struct MolBio l, 18 592–596. (DOI:10.1038/nsmb.2020).

[12] Kacar, B. & Gaucher, E. A. 2012 Towards the Recapitulation of Ancient History in the Laboratory: Combining Synthetic Biology with Experimental Evolution. In Artificial Life (pp. 11–18.

[13] Thornton, J. W. 2004 Resurrecting ancient genes: experimental analysis of extinct molecules. Nat Rev Genet, 5 366–375. (DOI:10.1038/nrg1324).

[14] Harms, M. J. & Thornton, J. W. 2013 Evolutionary biochemistry: revealing the historical and physical causes of protein properties. Nat Rev Genet, 14 559–571. (DOI:10.1038/nrg3540).

[15] Benner, S. A., Sassi, S. O. & Gaucher, E. A. 2007 Molecular paleoscience: systems biology from the past. Adv Enzymol Relat Areas Mol Biol 75, 1–132, xi.

[16] Koshi, J. M. & Goldstein, R. A. 1996 Probabilistic reconstruction of ancestral protein sequences. J Mol Evol, 42 313–320.

[17] Kacar, B., Ge, X., Sanyal, S. & Gaucher, E. A. 2017 Experimental Evolution of Escherichia coli Harboring an Ancient Translation Protein. J MolEvol. (DOI:10.1007/s00239–017-9781–0).

[18] Kacar, B. & Gaucher, E. A. 2012 Towards the Recapitulation of Ancient History in the Laboratory: Combining Synthetic Biology with Experimental Evolution. Artificial Life 13, 11–18.

[19] Whelan, S., Lio, P. & Goldman, N. 2001 Molecular phylogenetics: state-of-the-art methods for looking into the past. Trends in genetics : TIG 17, 262–272.

[20] Felsenstein, J. 2003 Inferring Phylogenies. MA.

[21] Yang, Z. & Rannala, B. 2012 Molecular phylogenetics: principles and practice. Nat Rev Genet, 13 303–314. (DOI:10.1038/nrg3186).

[22] Anisimova, M. 2012 Evolutionary Genomics: Statistical and Computational Methods. (New York, Humana Press.

[23] Risso, V. A., Gavira, J. A., Gaucher, E. A. & Sanchez-Ruiz, J. M. 2014 Phenotypic comparisons of consensus variants versus laboratory resurrections of Precambrian proteins. Proteins, 82 887–896. (DOI:10.1002/prot.24575).

[24] Sanderson, M. J. & Donoghue, M. J. 1996 Reconstructing shifts in diversification rates on phylogenetic trees. Trends Ecol Evol 11, 15–20.

[25] Williams, P. D., Pollock, D. D., Blackburne, B. P. & Goldstein, R. A. 2006 Assessing the accuracy of ancestral protein reconstruction methods. PLoS Comput Bio l, 2 e69. (DOI:10.1371/journal.pcbi.0020069).

[26] Cunningham, C. W., Omland, K. E. & Oakley, T. H. 1998 Reconstructing ancestral character states: a critical reappraisal. Trends Ecol Evol 13, 361–366.

[27] Loytynoja, A. & Goldman, N. 2008 Phylogeny-aware gap placement prevents errors in sequence alignment and evolutionary analysis. Science, 320 1632–1635. (DOI:10.1126/science.1158395).

[28] Adami, C. 2002 Sequence complexity in Darwinian evolution. Complexity 8, 59–56.

[29] Adami, C. 2004 Information theory in molecular biology. Physics of Life Reviews 1, 3–22.

[30] Jacobson, M. P., Friesner, R. A., Xiang, Z. & Honig, B. 2002 On the role of the crystal environment in determining protein side-chain conformations. Journal of molecular biology 320, 597–608.

[31] Jacobson, M. P., Pincus, D. L., Rapp, C. S., Day, T. J., Honig, B., Shaw, D. E. & Friesner, R. A. 2004 A hierarchical approach to all-atom protein loop prediction. Proteins, 55 351–367. (DOI:10.1002/prot.10613).

[32] Friesner, R. A., Murphy, R. B., Repasky, M. P., Frye, L. L., Greenwood, J. R., Halgren, T. A., Sanschagrin, P. C. & Mainz, D. T. 2006 Extra precision glide: docking and scoring incorporating a model of hydrophobic enclosure for protein-ligand complexes. Journal of medicinal chemistry, 49 6177–6196. (DOI:10.1021/jm051256o).

[33] Bershtein, S., Serohijos, A. W. & Shakhnovich, E. I. 2016 Bridging the physical scales in evolutionary biology: from protein sequence space to fitness of organisms and populations. Curr Opin Struct Biol 42, 31–40. (DOI:10.1016/j.sbi.2016.10.013).

[34] Dalziel, A. C., Rogers, S. M. & Schulte, P. M. 2009 Linking genotypes to phenotypes and fitness: how mechanistic biology can inform molecular ecology. Mol Ecol 18, 4997–5017. (DOI:10.1111/j.1365–294X.2009.04427.x).

[35] Dean, A. M. & Thornton, J. W. 2007 Mechanistic approaches to the study of evolution: the functional synthesis. Nat Rev Genet 8, 675–688. (DOI:10.1038/nrg2160).

[36] Eick, G. N., Bridgham, J. T., Anderson, D. P., Harms, M. J. & Thornton, J. W. 2016 Robustness of Reconstructed Ancestral Protein Functions to Statistical Uncertainty. Mol Biol Evol. (DOI:10.1093/molbev/msw223).

[37] Kacar, B. & Gaucher, E. A. 2013 Experimental evolution of protein-protein interaction networks. Biochem J 453, 311–319. (DOI:10.1042/BJ20130205).

[38] Lio, P. & Goldman, N. 1998 Models of Molecular Evolution and Phylogeny. Genome Res, 8 1233–1244.

[39] Yang, Z., Nielsen, R. & Goldman, N. 2000 Codon-substitution models for heterogeneous selection pressure at amino acid sites. Genetics 155, 421–449.

[40] Hanson-Smith, V., Kolaczkowski, B. & Thornton, J. W. 2010 Robustness of ancestral sequence reconstruction to phylogenetic uncertainty. Mol Biol Evol 27, 1988–1999. (DOI:10.1093/molbev/msq081).

[41] Altschul, S. F., Gish, W., Miller, W., Myers, E. W. & Lipman, D. J. 1990 Basic local alignment search tool. J Mol Biol215, 403–410. (DOI:10.1016/S0022–2836(05)80360–2).

[42] Anisimova, M., Liberles, D. A., Philippe, H., Provan, J., Pupko, T. & von Haeseler, A. 2013 State-of the art methodologies dictate new standards for phylogenetic analysis. BMC Evol Biol 13, 161. (DOI:10.11 86/1471–2148–13–161).

[43] Koonin, E. V. 2003 Comparative genomics, minimal gene-sets and the last universal common ancestor. Nature reviews. Microbiology 1, 127–136. (DOI:10.1038/nrmicro751).

[44] Mirkin, B. G., Fenner, T. I., Galperin, M. Y. & Koonin, E. V. 2003 Algorithms for computing parsimonious evolutionary scenarios for genome evolution, the last universal common ancestor and dominance of horizontal gene transfer in the evolution of prokaryotes. BMC Evol Biol 3, 2.

[45] Mushegian, A. 2008 Gene content of LUCA, the last universal common ancestor. Frontiers in bioscience : a journal and virtual library 13, 4657–4666.

[46] Benner, S. A., Ellington, A. D. & Tauer, A. 1989 Modern metabolism as a palimpsest of the RNA world. Proc Natl Acad Sci U S A 86, 7054–7058.

[47] Boller, A. J., Thomas, P. J., Cavanaugh, C. M. & Scott, K. M. 2011 Low stable carbon isotope fractionation by coccolithophore RubisCO. Geochim Cosmochim Ac 75, 7200–7207. (DOI:10.1016/j.gca.2011.08.031).

[48] Schopf, W. J. 2011 The paleobiological record of photosynthesis. Photosynth Res 107, 87–101. (DOI:10.1007/s 11120010–9577-1).

[49] Boller, A. J., Thomas, P. J., Cavanaugh, C. M. & Scott, K. M. 2015 Isotopic discrimination and kinetic parameters of RubisCO from the marine bloom-forming diatom, Skeletonema costatum. Geobiology 13, 33–43. (DOI:10.1111/gbi. 12112).

[50] McNevin, D. B., Badger, M. R., Whitney, S. M., von Caemmerer, S., Tcherkez, G. G. & Farquhar, G. D. 2007 Differences in carbon isotope discrimination of three variants of D-ribulose-1,5-bisphosphate carboxylase/oxygenase reflect differences in their catalytic mechanisms. J Biol Chem 282, 36068–36076. (DOI:10.1074/jbc.M706274200).

[51] Tcherkez, G. G., Farquhar, G. D. & Andrews, T. J. 2006 Despite slow catalysis and confused substrate specificity, all ribulose bisphosphate carboxylases may be nearly perfectly optimized. Proc Natl Acad Sci U S A 103, 7246–7251. (DOI:10.1073/pnas.0600605103).

[52] Tcherkez, G. G., Farquhar, G. D., Badeck, F. & Ghashgaie, J. 2004 Theoretical considerations about carbon isotope distribution in glucose of C3 plants. Functional Plant Biology 31, 857–877.

[53] Blankenship, R. E. & Hartman, H. 1998 The origin and evolution of oxygenic photosynthesis. Trends Biochem Sci, 23 94–97.

[54] Estep, M. F., Tabita, F. R., Parker, P. L. & Van Baalen, C. 1978 Carbon isotope fractionation by ribulose-1,5-bisophosphate carboxylase from various organisms. Plant physiology 61, 680–687.

[55] Tabita, F. R., Hanson, T. E., Li, H., Satagopan, S., Singh, J. & Chan, S. 2007 Function, structure, and evolution of the RubisCO-like proteins and their RubisCO homologs. Microbiol Mol Biol Rev, 71 576–599. (DOI:10.1128/MMBR.00015–07).

[56] Farquhar, G. D., Ehleringer, J. R. & Hubick, K. T. 1989 Carbon isotope discrimination and photosynthesis. Annu Rev Plant Bio 40, 503–537.

[57] Kacar, B., Hanson-Smith, V., Adam, Z. R. & Boekelheide, N. 2017 Constraining the timing of the Great Oxidation Event within the Rubisco Phylogenetic Tree. Geobiology.

[58] Shih, P. M., Occhialini, A., Cameron, J. C., Andralojc, P. J., Parry, M. A. & Kerfeld, C. A. 2016 Biochemical characterization of predicted Precambrian RuBisCO. Nat Commun 7, 10382. (DOI:10.1038/ncomms10382).

[59] Kacar, B., Adam, Z. R., Hanson-Smith, V. & Boekelheide, N. 2016 Constraining the Great Oxidation Event within the Rubisco phylogenetic tree. Geological Society of America 48, 289–213.

[60] Nisbet, E. G. & Nisbet, R. E. 2008 Methane, oxygen, photosynthesis, rubisco and the regulation of the air through time. Philos Trans R Soc Lond B Biol Sci, 363 2745–2754. (DOI:10.1098/rstb.2008.0057).

[61] Mangan, N. & Brenner, M. 2014 Systems analysis of the CO2 concentrating mechanism in cyanobacteria. eLife, e02043. (DOI:10.7554/eLife.02043).

[62] Badger, M. R. & Price, G. D. 2003 CO2 concentrating mechanisms in cyanobacteria: molecular components, their diversity and evolution. J Exp Bot 54, 609–622.

[63] Klughammer, B., Sultemeyer, D., Badger, M. R. & Price, G. D. 1999 The involvement of NAD(P)H dehydrogenase subunits, NdhD3 and NdhF3, in high-affinity CO2 uptake in Synechococcus sp. PCC7002 gives evidence for multiple NDH-1 complexes with specific roles in cyanobacteria. Mol Microbiol, 32 1305–1315.

[64] Kaplan, A. & Reinhold, L. 1999 Co2 Concentrating Mechanisms in Photosynthetic Microorganisms. Annu Rev Plant Physiol Plant Mol Biol 50, 539–570. (DOI:10.1146/annurev.arplant.50.1.539).

[65] Henry, R. P. 1996 Multiple roles of carbonic anhydrase in cellular transport and metabolism. Annual review of physiology 58, 523–538. (DOI:10.1146/annurev.ph.58.030196.002515).

[66] Hagemann, M., Kern, R., Maurino, V. G., Hanson, D. T., Weber, A. P. M., Sage, R. F. & Bauwe, H. 2016 Evolution of photorespiration from cyanobacteria to land plants, considering protein phylogenies and acquisition of carbon concentrating mechanisms. Journal of Experimental Botany 67, 2963–2976.

[67] Cousins, A. B., Badger, M. R. & von Caemmerer, S. 2006 Carbonic anhydrase and its influence on carbon isotope discrimination during C4 photosynthesis. Insights from antisense RNA in Flaveria bidentis. Plant physiology 141, 232–242. (DOI:10.1104/pp.106.077776).

[68] Lindskog, S. 1997 Structure and mechanism of carbonic anhydrase. Pharmacology & Therapeutics, 74 1–20.

[69] DeVoe, H. & Kistiakowsky, G. B. 1961 The Enzymic Kinetics of Carbonic Anhydrase from Bovine and Human Erythrocytes. J Am Chem Soc 83, 274–280.

[70] Smith, K. S. & Ferry, J. G. 2000 Prokaryotic carbonic anhydrases. FEMS Microbiol Rev 24, 335–366.

[71] Capasso, C. & Supura, C. T. 2014 An overview of the alpha-, beta- and gamma-carbonic anhydrases from Bacteria: can bacterial carbonic anhydrases shed new light on evolution of bacteria? Journal of Enzyme Inhibition and Medicinal Chemistry 30, 325–332.

[72] Hewett-Emmett, D. & Tashian, R. E. 1996 Functional diversity, conservation, and convergence in the evolution of the alpha-, beta-, and gamma-carbonic anhydrase gene families. Molecular phylogenetics and evolution 5, 50–77. (DOI:10.1 006/mpev. 1996.0006).

[73] Smith, K. S., Jacubzick, C., Whittam, T. S. & Ferry, J. G. 1999 Carbonic anhydrase is an ancient enzyme widespread in prokaryotes. Proc Natl Acad Sci U S A 96, 15184–15189.

[74] Banerjee, S. & Deshpande, P. A. 2016 On origin and evolution of carbonic anhydrase isozymes: A phylogenetic analysis from whole-enzyme to active site. Computational biology and chemistry 61, 121–129. (DOI:10.1016/j.compbiolchem.2016.01.003).

[75] Aizawa, K. & Miyachi, S. 1986 Carbonic anhydrase and CO2 concentrating mechanisms in microalgae and cyanobacteria. FEMS Microbiology Letters 39, 215–233.

[76] Graham, D., Reed, M. L., Patterson, B. D., Hockley, D. G. & Dwyer, M. R. 1984 Chemical properties, distribution, and physiology of plant and algal carbonic anhydrases. Annals of the New York Academy of Sciences 429, 222–237.

[77] Kumar, R. S. & Ferry, J. G. 2014 Prokaryotic carbonic anhydrases of Earth’s environment. Subcell Biochem 75, 77–87. (DOI:10.1007/978–94-007–7359-2_5).

[78] Tripp, B. C., Smith, K. & Ferry, J. G. 2001 Carbonic anhydrase: new insights for an ancient enzyme. J Biol Chem 276, 48615–48618. (DOI:10.1074/jbc.R100045200).

[79] Smeulders, M. J., Barends, T. R., Pol, A., Scherer, A., Zandvoort, M. H., Udvarhelyi, A., Khadem, A. F., Menzel, A., Hermans, J., Shoeman, R. L., et al. 2011 Evolution of a new enzyme for carbon disulphide conversion by an acidothermophilic archaeon. Nature 478, 412–416. (DOI:10.1038/nature10464).

[80] Mossel, E. & Steel, M. 2005 How much can evolved characters tell us about the tree that generated them?. In Mathematics of evolution and phylogeny. (ed. O. Gascuel), pp. 384–412. Oxford, Oxford University Press.

[81] Studer, R. A., Christin, P. A., Williams, M. A. & Orengo, C. A. 2014 Stability-activity tradeoffs constrain the adaptive evolution of RubisCO. Proc Natl Acad Sci U S A 111, 2223–2228. (DOI:10.1073/pnas. 1310811111).

[82] Woese, C. R., Olsen, G. J., Ibba, M. & Soll, D. 2000 Aminoacyl-tRNA synthetases, the genetic code, and the evolutionary process. Microbiol Mol Biol Rev 64, 202–236.

[83] Cronk, J. D., Endrizzi, J. A., Cronk, M. R., O’Neill J, W. & Zhang, K. Y. 2001 Crystal structure of E. coli beta-carbonic anhydrase, an enzyme with an unusual pH-dependent activity. Protein science : a publication of the Protein Society 10, 911–922. (DOI:10.1110/ps.46301).

[84] Rowlett, R. S. 2010 Structure and catalytic mechanism of the beta-carbonic anhydrases. Biochimica et biophysica acta 1804, 362–373. (DOI:10.1016/j.bbapap.2009.08.002).

[85] Khersonsky, O. & Tawfik, D. S. 2010 Enzyme promiscuity: a mechanistic and evolutionary perspective. Annual review of biochemistry 79, 471–505. (DOI:10.1146/annurev-biochem-030409–143718).

[86] Tawfik, D. S. 2010 Messy biology and the origins of evolutionary innovations. Nat Chem Biol, 6 692–696. (DOI:10.1038/nchembio.441).

[87] Bar-Even, A., Noor, E., Savir, Y., Liebermeister, W., Davidi, D., Tawfik, D. S. & Milo, R. 2011 The moderately efficient enzyme: evolutionary and physicochemical trends shaping enzyme parameters. Biochemistry 50, 4402–4410. (DOI:10.1021/bi2002289).

[88] Woese, C. R. 1967 The Genetic Code: The Molecular Basis for Genetic Expression. New York, Harper and Row.

[89] Aoshima, M. & Igarashi, Y. 2008 Nondecarboxylating and decarboxylating isocitrate dehydrogenases: oxalosuccinate reductase as an ancestral form of isocitrate dehydrogenase. J Bacteriol 190, 2050–2055. (DOI:10.1128/JB.01799–07).

[90] Aoshima, M., Ishii, M. & Igarashi, Y. 2004 A novel enzyme, citryl-CoA synthetase, catalysing the first step of the citrate cleavage reaction in Hydrogenobacter thermophilus TK-6. Mol Microbiol, 52 751–761. (DOI:10.1111/j.1365-2958.2004.04009.x).

[91] Lyons, T. W., Reinhard, C. T. & Planavsky, N. J. 2014 The rise of oxygen in Earth’s early ocean and atmosphere. Nature 506, 307–315. (DOI:10.1038/nature13068).

[92] Knoll, A. H. 2008 The Cyanobacteria: Molecular Biology, Genomics, and Evolution.,.

[93] Bell, E. A., Boehnke, P., Harrison, T. M. & Mao, W. L. 2015 Potentially biogenic carbon preserved in a 4.1 billion-year-old zircon. Proc Natl Acad Sci U S A 112, 14518–14521. (DOI:10.1073/pnas.1517557112).

[94] Schidlowski, M. 1988 A 3,800-Million-Year Isotopic Record of Life from Carbon in Sedimentary-Rocks. Nature 333, 313–318. (DOI:DOI 10.1038/333313a0).

[95] Grotzinger, J. P., Fike, D. A. & Fischer, W. W. 2011 Enigmatic origin of the largest-known carbon isotope excursion in Earth’s history. Nat Geoscience 4, 285–292.

[96] Gaucher, E. A., Govindarajan, S. & Ganesh, O. K. 2008 Palaeotemperature trend for Precambrian life inferred from resurrected proteins. Nature 451, 704–707. (DOI:10.1038/nature06510).

[97] Akanuma, S., Nakajima, Y., Yokobori, S., Kimura, M., Nemoto, N., Mase, T., Miyazono, K., Tanokura, M. & Yamagishi, A. 2013 Experimental evidence for the thermophilicity of ancestral life. Proc Natl Acad Sci U S A 110, 11067–11072. (DOI:10.1073/pnas.1308215110).

[98] Hobbs, J. K., Shepherd, C., Saul, D. J., Demetras, N. J., Haaning, S., Monk, C. R., Daniel, R. M. & Arcus, V. L. 2012 On the origin and evolution of thermophily: reconstruction of functional precambrian enzymes from ancestors of Bacillus. Mol BiolEvol, 29 825–835. (DOI:10.1093/molbev/msr253).

[99] Risso, V. A., Gavira, J. A., Mejia-Carmona, D. F., Gaucher, E. A. & Sanchez-Ruiz, J. M. 2013 Hyperstability and substrate promiscuity in laboratory resurrections of Precambrian beta-lactamases. J Am Chem Soc 135, 2899–2902. (DOI:10.1021/ja311630a).

[100] Woese, C. R. & Fox, G. E. 1977 The concept of cellular evolution. J Mol Evol 10, 1–6.

[101] Woese, C. R. 2002 On the evolution of cells. Proc Natl Acad Sci U S A 99, 8742–8747. (DOI:10.1073/pnas. 132266999).

[102] Woese, C. R. 1987 Bacterial evolution. Microbiological reviews 51, 221–271.

[103] Woese, C. 1998 The universal ancestor. Proc Natl Acad Sci U S A 95, 6854–6859.

[104] Fogel, M. L. & Cifuentes, L. A. 1993 Isotope fractionation during primary production. In Organic Geochemistry (eds. M. H. Engel & S. A. Macko). New Yok, Plenum.

[105] Finn, R. D., Clements, J., Arndt, W., Miller, B. L., Wheeler, T. J., Schreiber, F., Bateman, A. & Eddy, S. R. 2015 HMMER web server: 2015 update. Nucleic acids research, 43 W30–38. (DOI:10.1093/nar/gkv397).

[106] Katoh, K. & Standley, D. M. 2013 MAFFT multiple sequence alignment software version 7: improvements in performance and usability. Mol Biol Evol, 30 772–780. (DOI:10.1093/molbev/mst010).

[107] Capella-Gutierrez, S., Silla-Martinez, J. M. & Gabaldon, T. 2009 trimAl: a tool for automated alignment trimming in large-scale phylogenetic analyses. Bioinformatics, 25 1972–1973. (DOI:10.1093/bioinformatics/btp348).

[108] Stamatakis, A. 2014 RAxML version 8: a tool for phylogenetic analysis and post-analysis of large phylogenies. Bioinformatics 30, 1312–1313.

[109] Hanson-Smith, V. & Johnson, A. 2016 PhyloBot: A Web Portal for Automated Phylogenetics, Ancestral Sequence Reconstruction, and Exploration of Mutational Trajectories. PLoS Comput Biol 12, e1004976. (DOI:10.1371/journal.pcbi.1004976).

[110] Waterhouse, A. M., Procter, J. B., Martin, D. M., Clamp, M. & Barton, G. J. 2009 Jalview Version 2–a multiple sequence alignment editor and analysis workbench. Bioinformatics 25, 1189–1191. (DOI:10.1093/bioinformatics/btp033).

[111] Huang, S., Hainzl, T., Grundstrom, C., Forsman, C., Samuelsson, G., Sauer-Eriksson, A.E. Structural Studies of β-Carbonic Anhydrase from the Green Alga Coccomyxa: Inhibitor Complexes with Anions and Acetazolamide. PLoS ONE. https://doi.org/10.1371/journal.pone.0028458

